# GABAergic inhibition of leg motoneurons is required for normal walking behavior in freely moving *Drosophila*

**DOI:** 10.1101/173120

**Authors:** Swetha B.M. Gowda, Pushkar D. Paranjpe, O. Venkateswara Reddy, Sudhir Palliyil, Heinrich Reichert, K. VijayRaghavan

**Affiliations:** National Centre for Biological Sciences, Tata Institute of Fundamental Research, Bangalore 560 065, India; Manipal University, Manipal 576 104, Karnataka, India; Biozentrum, University of Basel, Basel 4001, Switzerland.

**Keywords:** Walking, Pre-motor inhibition, intemeurons, leg motoneurons, Speed

## Abstract

Walking is a complex rhythmic locomotor behaviour generated by sequential and periodical contraction of muscles essential for coordinated control of movements of legs and leg joints. Studies of walking in vertebrates and invertebrates have revealed that premotor neural circuitry generates a basic rhythmic pattern that is sculpted by sensory feedback and ultimately controls the amplitude and phase of the motor output to leg muscles. However, the identity and functional roles of the premotor interneurons that directly control leg motoneuron activity are poorly understood. Here we take advantage of the powerful genetic methodology available in *Drosophila* to investigate the role of premotor inhibition in walking by genetically suppressing inhibitory input to leg motoneurons. For this, we have developed a novel algorithm for automated analysis of leg motion to characterize the walking parameters of wildtype flies from high speed video recordings. Further, we use genetic reagents for targeted RNAi knockdown of inhibitory neurotransmitter receptors in leg motoneurons together with quantitative analysis of resulting changes in leg movement parameters in freely walking *Drosophila*. Our findings indicate that targeted down regulation of GABAA receptor Rdl in leg motoneurons results in a dramatic reduction of walking speed and step-length without the loss of general leg coordination during locomotion. Genetically restricting the knockdown to the adult stage and subsets of motoneurons yields qualitatively identical results. Taken together, these findings identify GABAergic premotor inhibition of motoneurons as an important determinant of correctly coordinated leg movements and speed of walking in freely behaving *Drosophila*.

**SIGNIFICANCE STATEMENT:** Inhibition is an important feature of neuronal circuit and in walking it aids in controlling coordinated movement of legs, leg segments and leg joints. Recent studies in *Drosophila* reports the role of premotor inhibitory interneurons in regulation of larval locomotion. However, in adult walking the identity and functional role of premotor interneurons is less understood. Here, we use genetic methods for targeted knockdown of inhibitory neurotransmitter receptor in leg motoneurons that results in slower walking speed and defects in walking parameters combined with novel method we have developed for quantitative analysis of the fly leg movement and the observed changes in walking parameters. Our results indicate that GABAergic pre-motor inhibition to leg motoneurons is required to control the normal walking behaviour in adult *Drosophila*.

## INTRODUCTION

Walking is a complex rhythmic locomotor behavior that requires the coordinated control of movements among legs, leg segments and leg joints (1–5).While the complete neural circuitry of the motor control networks that orchestrate this control is not known in any animal, significant progress has been made in understanding the general organization of the underlying networks. Studies of reduced preparations in vertebrates have revealed that premotor interneuronal circuits present in the spinal cord are capable of generating patterned rhythmic locomotor activity and distributing this activity to motoneurons (6–8). Furthermore, in several cases the function of identified spinal interneuron types in this type of fictive locomotion has been elucidated (9, 10). However, the complexity and the difficulty of genetic intervention in vertebrate models have made the cellular identification and functional analysis of these premotor control circuits difficult.

Invertebrate models have reduced complexity and for some of them, such as *Drosophila*, genetic reagents that enable visualization and perturbation of neural circuitry are available (11–13). In *Drosophila*, the neural connectivity of leg motoneurons, their structural organization and their relationship with leg muscles has been well established (14–17). Moreover, extensive neurogenetic analyses of walking behavior in *Drosophila* have shown that higher brain centers such as central complex, the ellipsoid body and the mushroom bodies affect higher order phenomenon such as the drive to walk, ability to maintain a fixed course or goal directed locomotion (18,19,21). However, these central brain structures are entirely dispensable for the more mundane aspect of walking – maintaining all the moving parts in harmony with each other.

The observation that headless flies can walk, demonstrates that the higher order brain regions has less contribution for coordination of leg motion and that the minimal circuitry required for the generation of rhythmic walking behavior resides in the ventral nerve cord (22,23). Thus, as in other insects, premotor neural circuitry in the ventral nerve cord is thought to generate a basic rhythm, which is sculpted by bilateral and intersegmental coordination processes; modified by sensory feedback such that it ultimately controls the magnitude and timing of patterned motoneuronal output to leg muscles as the animal walks (20,24–26).

Inhibitory interactions are important features of the neural circuitry that generate walking (27-29). Inhibition is involved in controlling flexor-extensor alternation, in bilateral and intersegmental coordination as well as in propagation and termination of motoneuron activity as shown in vertebrates (9, 30, 31). In *Drosophila*, recent studies on larval stages have identified two types of inhibitory premotor interneurons that are involved in controlling the motor activity required for larval crawling (32, 33). Moreover, also in larvae, segmentally repeated GABAergic interneurons have been identified and implicated in the control of the peristaltic wave of activity that underlies larval crawling (34). However, in adult flies the identity and functional circuit features of the premotor interneurons that control leg motoneuron activity are poorly understood and virtually nothing is known about a role of inhibitory premotor interneurons in walking behavior.

Here we investigate the role of premotor inhibition in walking by selectively suppressing GABAergic input to leg motoneurons. For this, we first develop a novel algorithm for the automated analysis of leg movements in order to characterize the walking parameters of wild type flies from high speed video recordings. Using genetic reagents that allow selective labeling of leg motoneurons together with targeted RNAi knockdown of neurotransmitter receptors, we then interrogate the nature of their pre-motor neuronal input during walking by analyzing the walking parameters. Our findings indicate that knocking-down the expression of GABA_A_ receptor Rdl (Resistance to dieldrin) results in dramatically reduced walking speed as well as reduced step-length and failure to achieve sustained leg extensions during locomotion. Genetically restricting the knockdown to the adult stage or to subsets of motoneurons gives qualitatively identical results - slower walking speeds, shortened step lengths without a general loss of coordination. Altogether, these findings identify premotor inhibition of motoneurons as an important determinant of leg movement and resulting speed in freely behaving *Drosophila*.

## RESULTS

### Recording and analysis of leg movements in freely walking flies

As a prerequisite for investigating whether pre-motor inhibitory inputs are required for appropriate walking behavior in *Drosophila*, we developed an automated high-speed video recording and analysis technique. This technique made it possible to record the protraction (swing) and retraction (stance) phases of leg motion at high resolution for each leg in freely walking flies **(see methods)**. For a detailed characterization of these leg movement phases, an auto-detection of leg movements from video frames was carried out followed by the quantitative analysis of leg movements from the extracted leg-tip coordinate data.

To auto-detect the movement of all six legs in their entirety given the video frame of a fly, a novel algorithm was developed using an open source image analysis software package - Fiji (2013 lifeline version). In this algorithm, light intensity thresholding was used to obtain a binary image of the fly and its contour was auto-selected as a full ROI (region of interest). Subsequently, the full ROI was adjusted such that legs were uncovered (torso ROI) and individual leg ROIs are computed by pixel-wise logical XOR between the full ROI and the torso ROI (Fig. 2A). (Alternatively, in simplified terms, this algorithm can be described according to binary morphological operations of erode and dilate as follows: erode the binary mask of the fly n times such that only the partially eroded torso remains, dilate the binary mask of the torso n times in order to restore the size of the torso, compute a third binary mask by performing a pixel-wise XOR operation between the full fly binary mask and the torso binary mask, obtain the binary mask of all six legs without the torso). After the leg’s ROIs were isolated, the leg-tip coordinates were extracted for further analysis.

**Figure 1:**
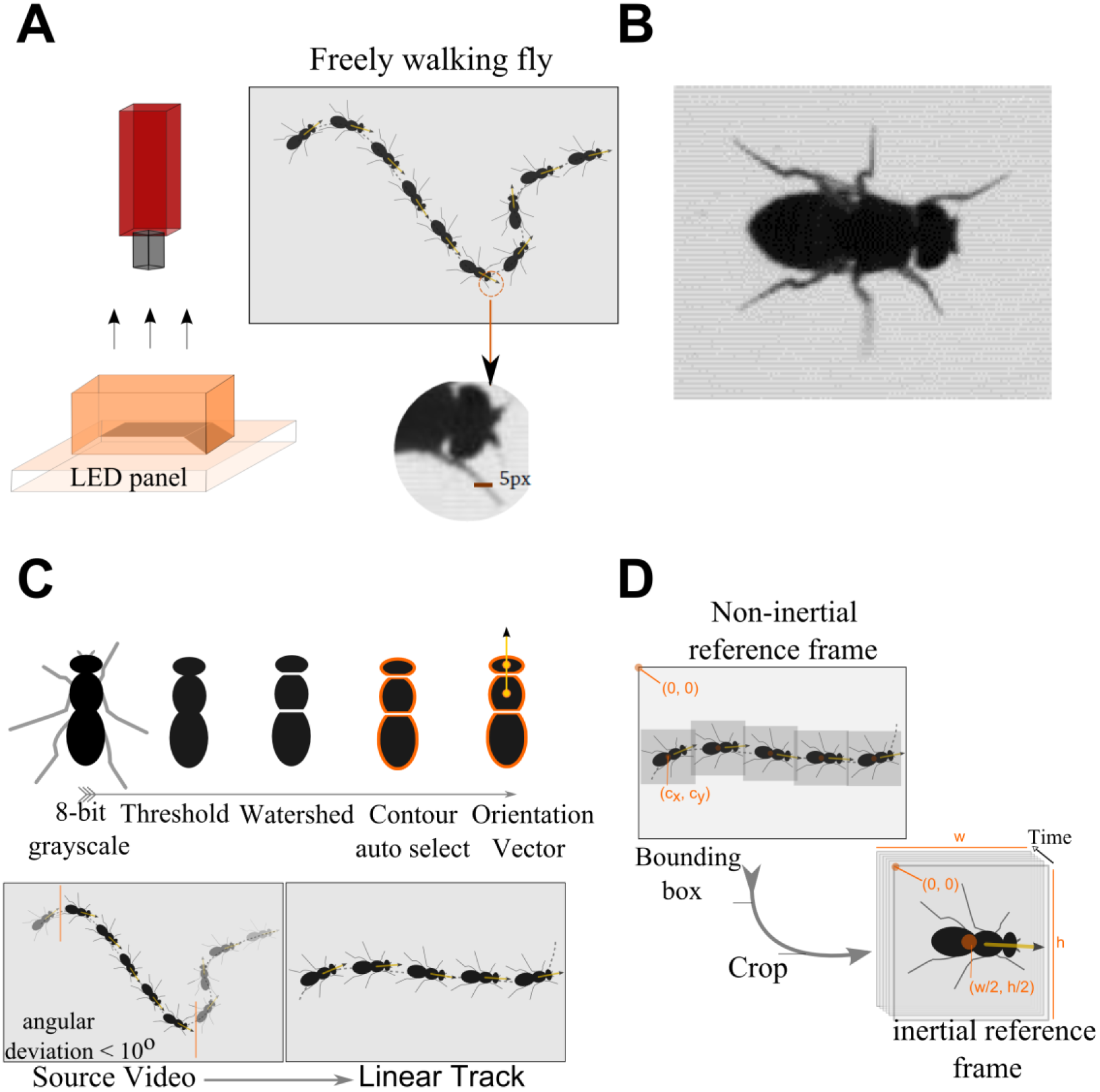
Automated recording of walking behaviour of freely moving flies. (A)Leg movements of freely walking flies were recorded with a 200fps camera; flies were placed in the rectangular shaped arena illuminated from below with LED light source. (B) Dorsal view of the fly captured at high resolution. (C) To obtain linear walking sequences, each frame is thresholded such that only the torso is visible. A watershed filter cuts the torso into three sections - the head, the thorax and the abdomen. The centroids of the head and the body define the orientation vector. Enforcing a maximum angular deviation of 10° as a chief criterion, contiguous frames that constitute a linear walking bout are selected. (D) A minimum bounding box that encloses the fly body is used as a mask to crop each frame and rotational transformation is applied so that the fly’s linear motion occurs along the positive direction of the new X-axis. The centroid of the fly was computed in each frame and the origin of the new coordinated axis was shifted to centroid. This transforms the non-inertial frame of reference to inertial frame of reference such that the fly appears to be walking on a treadmill.

**Figure 2:**
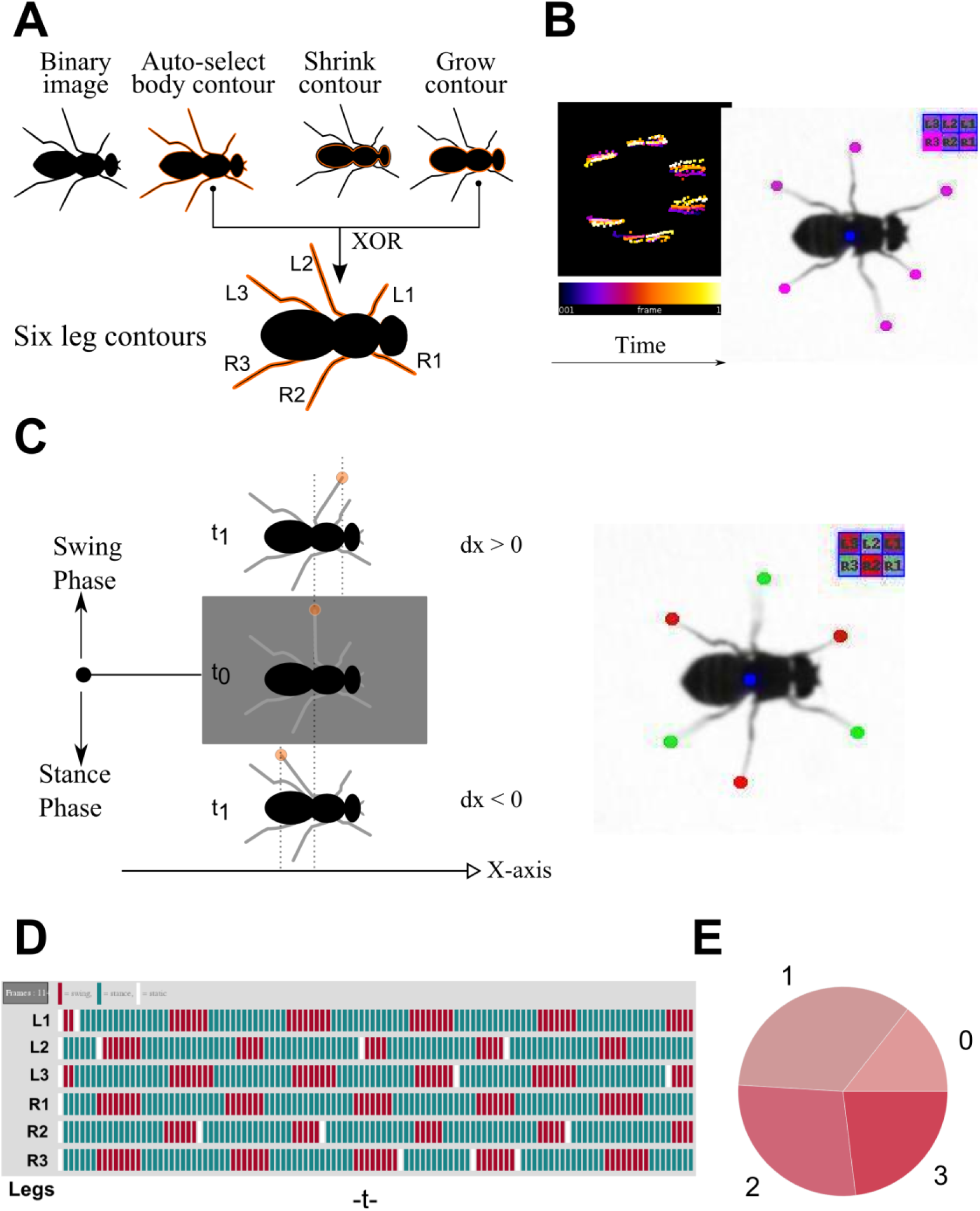
Automated detection of leg movements in freely walking flies. (A) Fly body contour is auto-selected and shrunk by 5px.The contour selection is grown back by 5px as a result of which only the torso gets selected, leaving the legs out. The fly contour selection and the fly torso selection are combined through a XOR function. This gives a composite selection containing all six legs. (B) Temporal Z-projection of leg-tip trajectories allows rapid extraction of individual leg-tips by drawing region of interest selections around six clusters. The temporally color-coded heat-map is a record of the trajectory of the leg-tip during the entire walking bout and aids in demarcating adjacent legs. The leg-tips and the body centroid are annotated in every frame of the video in order to aid in assessing the quality of their detection. (C) Swing phase and stance phase classification is performed based on the sign of change in the X co-ordinate of the leg tip with respect to time. A positive change implies swing phase of the leg while negative change implies stance phase of the leg. A swinging leg was annotated by overlaying a green filled circle on its tip; a stancing leg was annotated by overlaying a red filled circle on its tip. No leg was annotated if it was stationary (D) The swing, stance or steady phase of each of the six legs with respect to time is represented as series of colored ticks in a gait diagram. A representative gait diagram is shown in which each tick is 5ms in duration; red tick implies swing, turquoise tick implies stance and white tick implies steady phase. (E) Three legs swinging concurrently defines a concurrency state of 3, two legs swinging concurrently defines a concurrency state of 2, one leg swinging alone defines a concurrency state of 1, while state 0 implies no leg was swinging.

A temporal projection of the leg-tip coordinates resulted in an image that consisted of six distinct clusters, which allowed efficient isolation and assignment of each leg-tip coordinate to the respective hemithoracic segment (Fig. 2B). Each leg-tip coordinate was a 3-dimensional entity comprising x, y and frame number values. From these values, two distinct phases of leg motion could be identified, namely the protraction (swing) phase in which the leg moves in a posterior to anterior direction and the retraction (stance) phase in which the leg moves in an anterior to posterior direction (2C). During the protraction (swing) phase the leg is raised from the substrate and is moving in a posterior to anterior direction relative to the body; during the retraction (stance) phase the leg is in contact with the substrate and is moving in an anterior to posterior direction relative to the body. The dynamics of swing/stance events for each of the six legs time was represented in a diagram similar to the “gait diagram” traditionally used to visualize walking gaits (Fig. 2D). To indicate the degree of coordination between legs, concurrency scores were calculated and depicted in the concurrency score pie chart (Fig. 2E).

### Characterization of swing and stance phases in freely walking wild type flies

In an initial baseline behavioral analysis of freely walking flies we used this technique to analyze the relationship between walking speed and leg protraction (swing) and retraction (stance) phase characteristics in the wild type. For this, we acquired approximately 39 video recordings of flies walking freely in an arena. These video recordings were grouped into two classes with respect to walking speed. These classes are fast (21.0 mm/s to 34.0 mm/s) and slow (7.0 mm/s to 20.0 mm/s) see **supplementary movie 1**.

Analysis of recorded leg movements indicated that faster walking flies protracted their legs farther (increased swing amplitude) than slower walking flies (Fig. 3A). Similarly, faster walking flies retracted their legs farther (increased stance amplitude) when compared to slower walking flies (Fig. 3C). The largest difference in the length of protraction between the fastest walking group and the slowest walking group was ~300um; this difference amounts to about one third of the average body length of the animal. Faster walking flies completed their leg protractions more rapidly (decreased swing duration) than the slower walking groups (Fig. 3B). Faster walking flies also completed their leg retractions more rapidly (decreased stance duration) than the slower walking groups (Fig. 3D).

**Figure 3:**
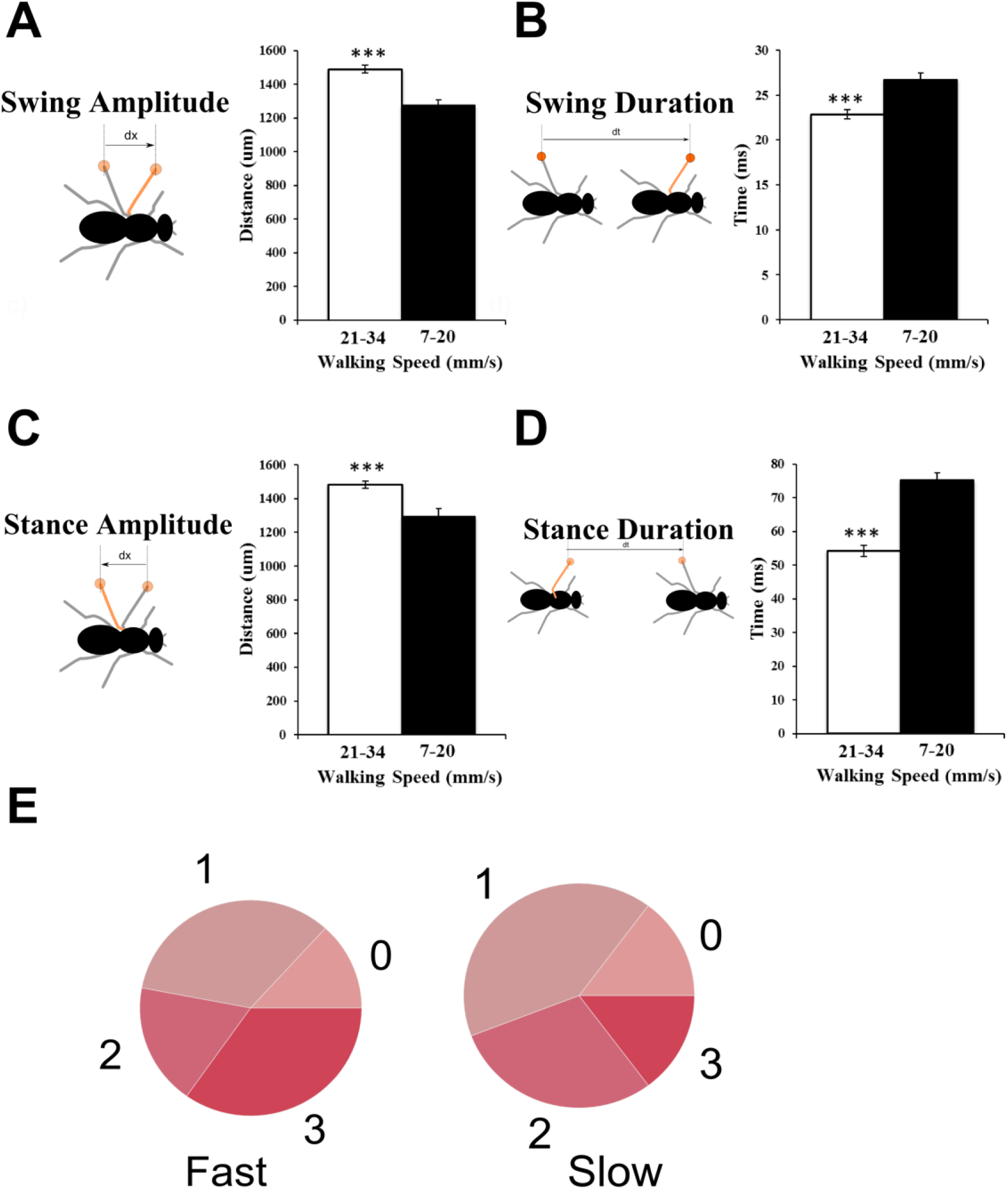
Relationship between walking speed and leg phase characteristics in freely walking flies. Leg movement parameters are quantified and classified into two speed intervals; slow (7.0-21.0 mm/s) and fast (21.0-34.0) mm/s.) (A) Swing amplitude is the average displacement of the leg-tip during swing phase as calculated in the fly’s inertial reference frame; it increases as the walking speed increases. (B) Swing duration is the time of the swing phase; it decreases as the walking speed increases. (C) Stance amplitude is the average displacement of the leg-tip during stance phase as calculated in the fly’s inertial reference frame; it increases as the walking speed increases. (D) Stance duration is the time stance phase; it decreases as the walking speed increases. (E) Concurrency state proportions to change with respect to walking speed. State 3 is proportionately higher at faster walking speeds but lower at slower walking speeds. N=39 videos. All the bar represents mean ± SEM (^∗∗∗^: P <= 0.001; ^∗∗^: P <= 0.01; ^∗^: P <= 0.05.). Student’s t-test was performed for all statistical analysis.

The largest difference in the duration of retraction (stance duration) between the fastest and the slowest walking group was ~30ms. The fastest walking animals thus complete each metachronal wave in ~33% less time than the slowest walking animal. We also observed that the concurrency state proportions change with respect to walking speed. Concurrency state 3 corresponding to a tripod gait is proportionately more prevalent for faster walking flies than for slower walking flies (Fig. 3E). Overall these results quantify the way in which wild type flies alter the length and duration of leg swing and stance phases as they modulate their walking speed.

### Targeted knockdown of Rdl provides evidence for GABAergic pre-motor inhibitory input to leg motoneurons during walking

To determine if pre-motor inhibition of leg motoneurons might be important for correct walking behavior, we conducted a screen for walking deficits caused by targeted knockdown of neurotransmitter receptors. For this, UAS-RNAi constructs for receptors of the neurotransmitters GABA, glutamate and acetylcholine were used together with the OK371Gal4 driver, which targets UAS transgene expression to glutamatergic neurons and hence labels all leg motoneurons (Fig. 4A). Expression of OK371Gal4 is controlled by the enhancer of the *Drosophila* vesicular glutamate transporter gene, VGLUT; (35).

**Figure 4:**
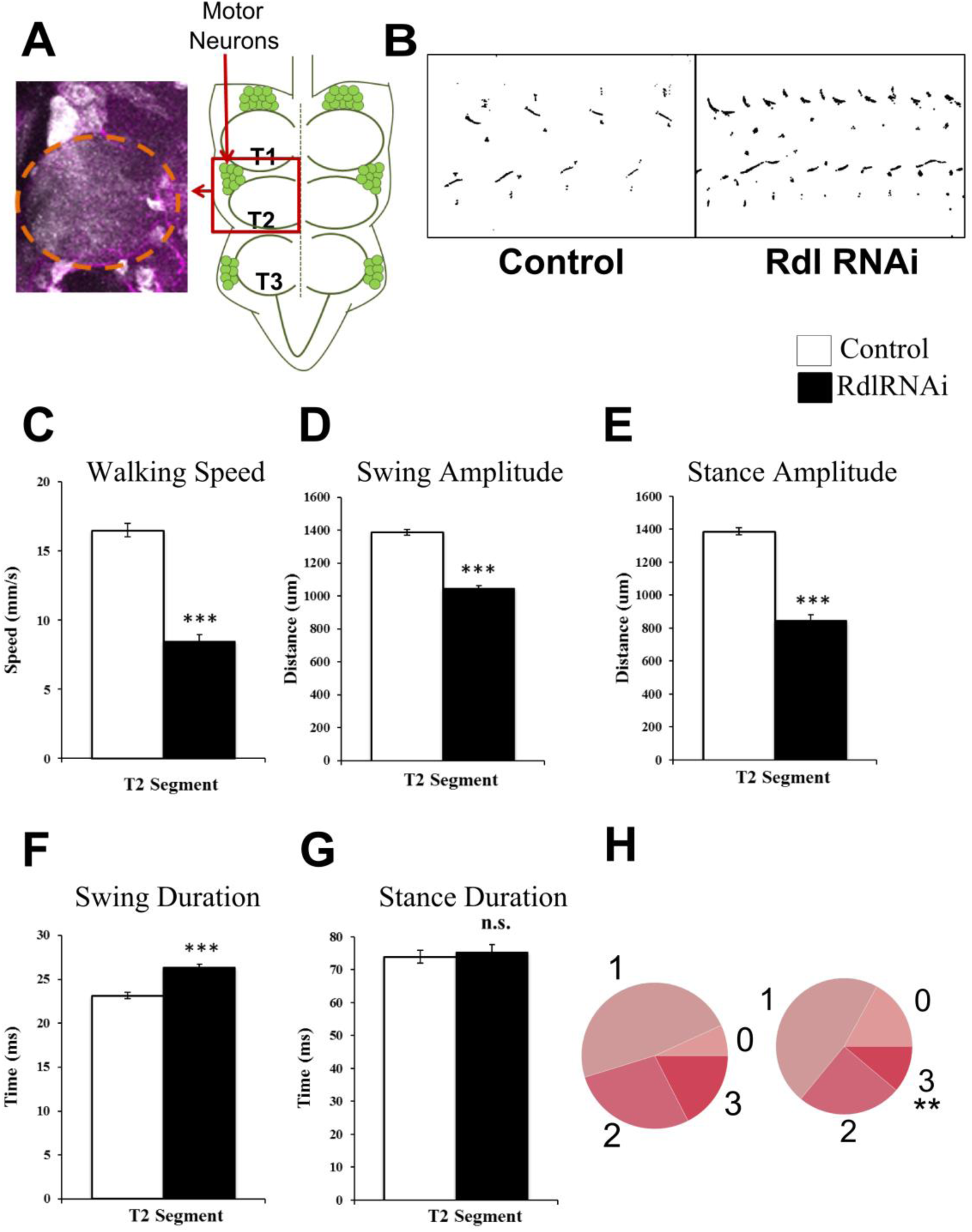
Targeted knockdown of Rdl provides evidence for GABAergic inhibitory input to motoneurons in walking flies. (A) The OK371-GAL4 driver targets UASmCD8GFP expression to leg motoneurons. Schematic representation of motoneuron cell bodies localized in the ventral nerve cord (right). Enlarged T2 hemineuromere region showing GFP marked motor neuron cell bodies and dendritic arborization (left). Orange circled region indicates neuropil. (B) Rdl receptor knockdown using OK371 GAL4 in all leg motoneurons results in shorter step length as revealed by the soot imprints. (C) Rdl receptor knockdown using OK371 GAL4 in all leg motoneurons results in significant decrease in walking speed. In this and all subsequent figures left bar indicates control, right bar indicates knockdown. D,E,F) Rdl receptor knockdown using OK371 GAL4 in all leg motoneurons results in reduced amplitudes of swing and stance and increased swing duration. g) Stance duration is unaffected. H) Concurrency state 3 is significantly decreased by Rdl receptor knockdown. Left circle control, right circle knockdown. Controls N=76, Experiment N=68. All the bar represents mean ± SEM (^∗∗∗^: P <= 0.001; ^∗∗^: P <= 0.01; ^∗^: P <= 0.05.). Student’s t-test was performed for all statistical analysis.

Initial screening for walking deficits in these RNAi knockdown experiments was carried out using a soot assay of leg footprints (36, 17). In this prescreen, the targeted knockdown of the GABA_A_ receptor Rdl (Resistance to dieldrin) resulted in a prominent walking phenotype characterized by shorter step length as compared to controls (Fig. 4B). For a more extensive characterization of the walking behavior deficits that resulted from this targeted Rdl knockdown, we used our automated video analysis to characterize the relationship between walking speed and leg swing and stance phase characteristics in a quantitative manner. Analysis of the video data revealed that the Rdl knockdown flies had a markedly reduced walking speed as compared to controls (Fig. 4C) see **supplementary movie 2**. Moreover, the amplitude of the leg swing phase and consequently that of leg stance phase was markedly shorter as compared to controls (Fig. 4D and 4E). Swing phase duration was significantly higher than controls (Fig. 4F) but there was no change in the duration of stance phases (Fig. 4G). The concurrency state 3, in which three legs swing nearly simultaneously, was proportionally reduced as compared to controls (Fig. 4H). Shorter average swing amplitudes in every metachronal walking wave markedly reduced the reach of the animal, thereby reducing the walking speed. (Data shown here is for T2 leg segment as a representative of other leg segments)

Given that Rdl is an ionotropic inhibitory neurotransmitter receptor, these results suggest that normal, wild type-like walking behavior requires inhibitory pre-motor input to leg motoneurons mediated by GABA acting through the Rdl receptor.

### Adult-specific targeted Rdl knockdown in leg motoneurons also results in abnormal walking behavior

Suppression of inhibitory pre-motor input by constitutive knockdown of Rdl throughout all developmental stages could potentially impact the formation of the walking circuit. To rule out the possibility that developmental effects were not responsible for the observed walking phenotypes, we refined our analysis by performing adult-specific knockdown of Rdl in leg motor neurons. For this we used the temperature sensitive Gal4 repressor, Gal80ts, together with OK371Gal4 to limit the effects of the targeted knockdown to adult stages. Flies were grown at 18°Cand shifted to 29°C post eclosion (Fig. 5A; Gal80ts is active at 18°Cand is inactive at 29°C).

**Figure 5:**
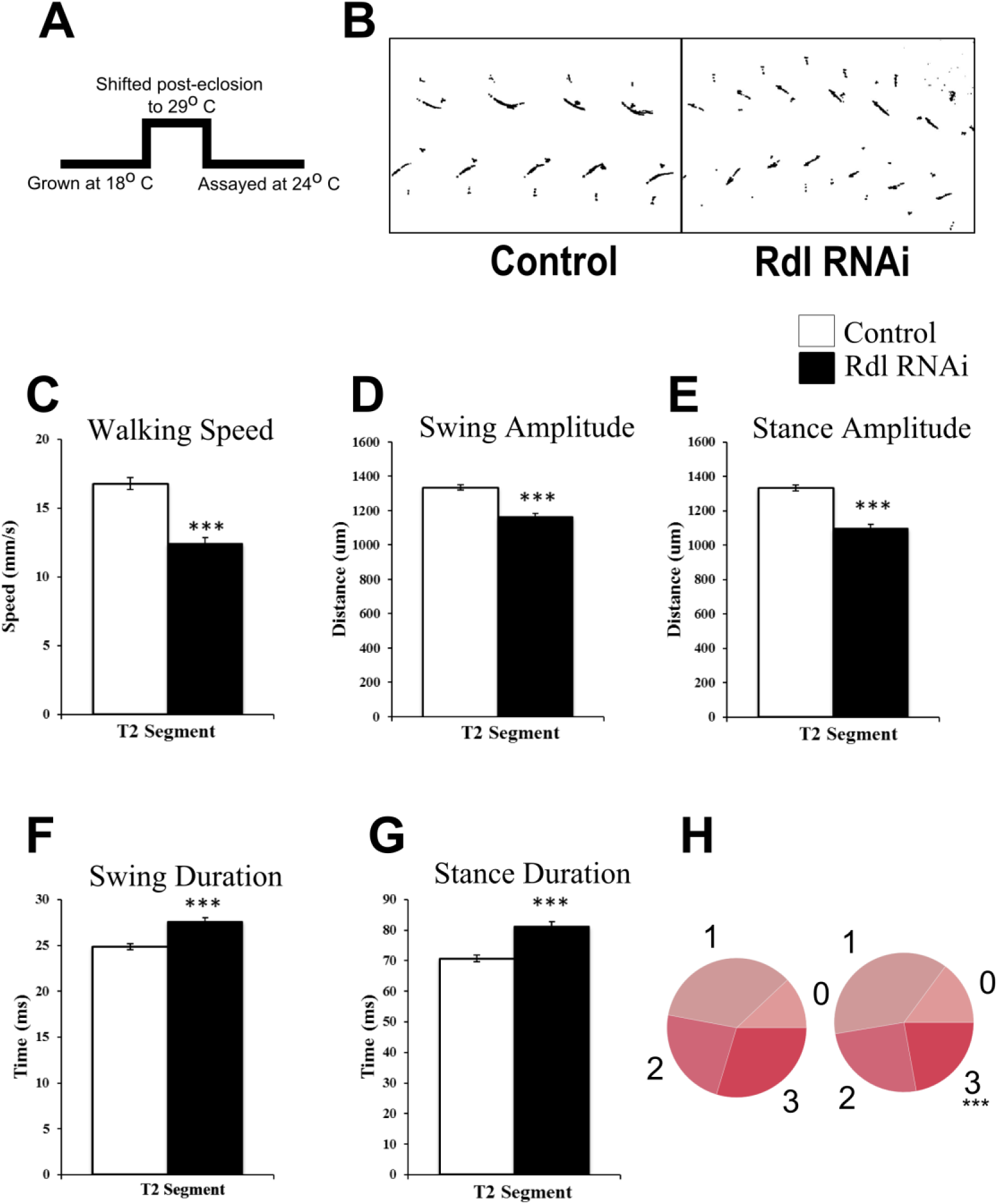
Adult-specific Rdl knockdown in leg motoneurons results in abnormal walking behaviour. (A) For adult specific knockdown of Rdl receptor, the GAL4 repressor Gal80ts was used to limit the effects to adult stages. For this flies were grown at 18°C and shifted to 29°C post eclosion. (B) Adult-specific Rdl receptor knockdown using OK371 GAL4 in all leg motoneurons results in shortened step length as shown in soot print images (C) Rdl receptor knockdown flies show slower walking speed. In this and all subsequent figures left bar indicates control, right bar indicates knockdown. (D,E) Amplitudes of swing and stance are shorter in Rdl knockdown flies as compared to controls.( F,G) Swing and stance duration are increased longer in Rdl knockdown flies as compared to controls. (H) Concurrency state 3 is significantly reduced in Rdl receptor knockdown flies. Left circle control, right circle knockdown. Controls N=108, Experiment N=77. All the bar represents mean ± SEM (^∗∗∗^: P <= 0.001; ^∗∗^: P <= 0.01; ^∗^: P <= 0.05.). Student’s t-test was performed for all statistical analysis.

Soot assay prints reveal that adult-specific targeted Rdl knockdown also caused walking behavior phenotypes characterized by shorter step length as compared to controls (Fig.5B). More detailed quantitative characterization of the walking phenotypes using video analysis showed that these flies had a markedly reduced walking speed as compared to controls (Fig. 5C) see **supplementary movie 3**. Moreover, the amplitude of the leg swing phase and consequently that of leg stance phase was markedly shorter as compared to controls (Fig. 5D and 5E) and the durations of swing and stance phases were significantly higher than in controls (Fig 5F and 5G). The concurrency state 3, in which three legs swing together (tripod gait), was proportionally reduced as compared to controls (Fig. 5H). (Data shown here is for T2 leg segment as a representative of other leg segments)

These results indicate that abnormal walking behavior also results if GABAergic inhibitory pre-motor input to leg motoneurons acting through the Rdl receptor is impaired specifically in mature adult stages

### Leg motoneuron subset-specific knockdown of Rdl also results in abnormal walking behavior

In the above-mentioned experiments, the inhibitory GABA_A_ receptor Rdl was down regulated in all of the fly’s motoneurons by the OK371 Gal4 driver. To determine if targeted knockdown of Rdl limited to only a subset of the leg motoneurons might also result in aberrant walking behavior, we took advantage of a generated VGN 1-Intron Reg-3 Gal4 driver. This driver targets reporter gene expression to only a small subset of leg motoneurons that innervate the trdm, fedm, tidm, and tadm muscles of the leg (Fig 6A).

**Figure 6:**
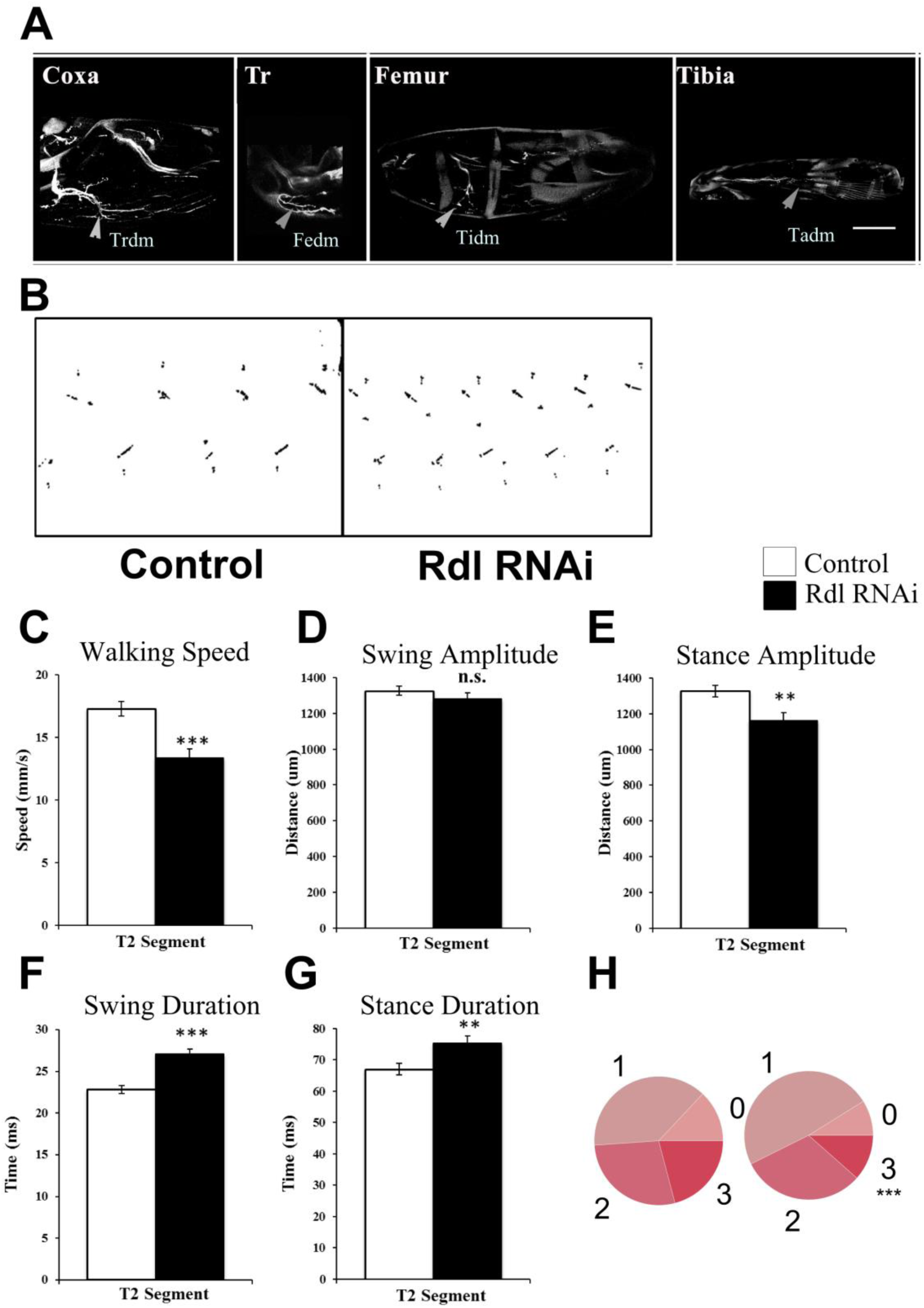
Subset specific knockdown of Rdl in leg motoneurons results in deficits in walking behaviour. A) Innervation patterns of leg motoneurons labeled by VGN 1-Intron Reg-3 GAL4 driving UAS-GFP (Scale bar 50 μm). Motoneuron axons innervating most of the depressor muscles (trdm-trochanter depressor muscle, fedm-femur depressor muscle, tidm-tibia depressor muscle, tadm-tarsal depressor muscle) are GFP labelled. B) Motoneuron subset specific Rdl receptor knockdown results in shortened step lengths as shown in the soot prints. C) Rdl knockdown in subsets of leg motoneurons results in reduced walking speed. In this and all subsequent figures left bar indicates control, right bar indicates knockdown. D,E) Rdl knockdown in subsets of leg motoneurons results in decreased stance amplitude; swing amplitude is not affected. F,G) Rdl knockdown in subsets of leg motoneurons results in longer swing duration and stance duration. H) Concurrency state 3 is significantly reduced in Rdl receptor knockdown flies. Left circle control, right circle knockdown. Controls N=48, Experiment N=45. All the bar represents mean ± SEM (^∗∗∗^: P <= 0.001; ^∗∗^: P <= 0.01; ^∗^: P <= 0.05.). Student’s t-test was performed for all statistical analysis.

Soot assay print assays reveal that motoneuron subset-specific targeted Rdl knockdown also caused walking behavior phenotypes (Fig 6B). Detailed quantitative characterization of these walking phenotypes using video analysis showed that the flies had reduced walking speeds (Fig 6C) see **supplementary movie 4**, and the amplitude of leg stance was markedly shorter; but no change in swing amplitude was observed (Fig. 6D-E). The durations of leg swing and stance phases were significantly different as compared to wildtype controls (Fig. 6F-G). The concurrency state 3, in which three legs protract nearly simultaneously, was proportionally reduced as compared to controls (Fig. 6H). The magnitude of the walking deficits caused by VGN 1-Intron Reg-3 Gal4 targeted Rdl knockdown in leg motoneuron subsets was generally smaller than those observed in OK371Gal4 targeted knockdown of Rdl in all leg motoneurons. (Data shown here is for T2 leg segment as a representative of other leg segments)

Nevertheless, these findings suggest that the suppression of premotor inhibitory input, even if it is limited to a small subset of leg motoneurons, is sufficient for perturbation of normal walking behavior.

### Analysis of Rdl knockdown in a Tsh-Gal80 background confirms leg motoneurons as sites of premotor inhibition

The behavioral phenotypes caused by OK371Gal4 targeted knockdown of Rdl indicate that reduced GABAergic inhibition in the neurons genetically accessed by this Gal4 driver (“OK371 neurons”), results in walking behavior deficits. Given that the leg motoneurons are prominent among the OK371 neurons, it seems likely that impairment of GABA-ergic inhibition of these motoneurons is responsible for the observed walking defects. However, the full complement of OK371 neurons as visualized by using OK371Gal4 to drive a UAS-GFP reporter includes numerous interneurons in the central brain and optic lobes, in addition to the leg motoneurons (Fig. 7A). Might impairment in the GABA-ergic inhibition of these central brain interneurons contribute to the walking defects observed in the OK371 targeted knockdown experiments?

**Figure 7:**
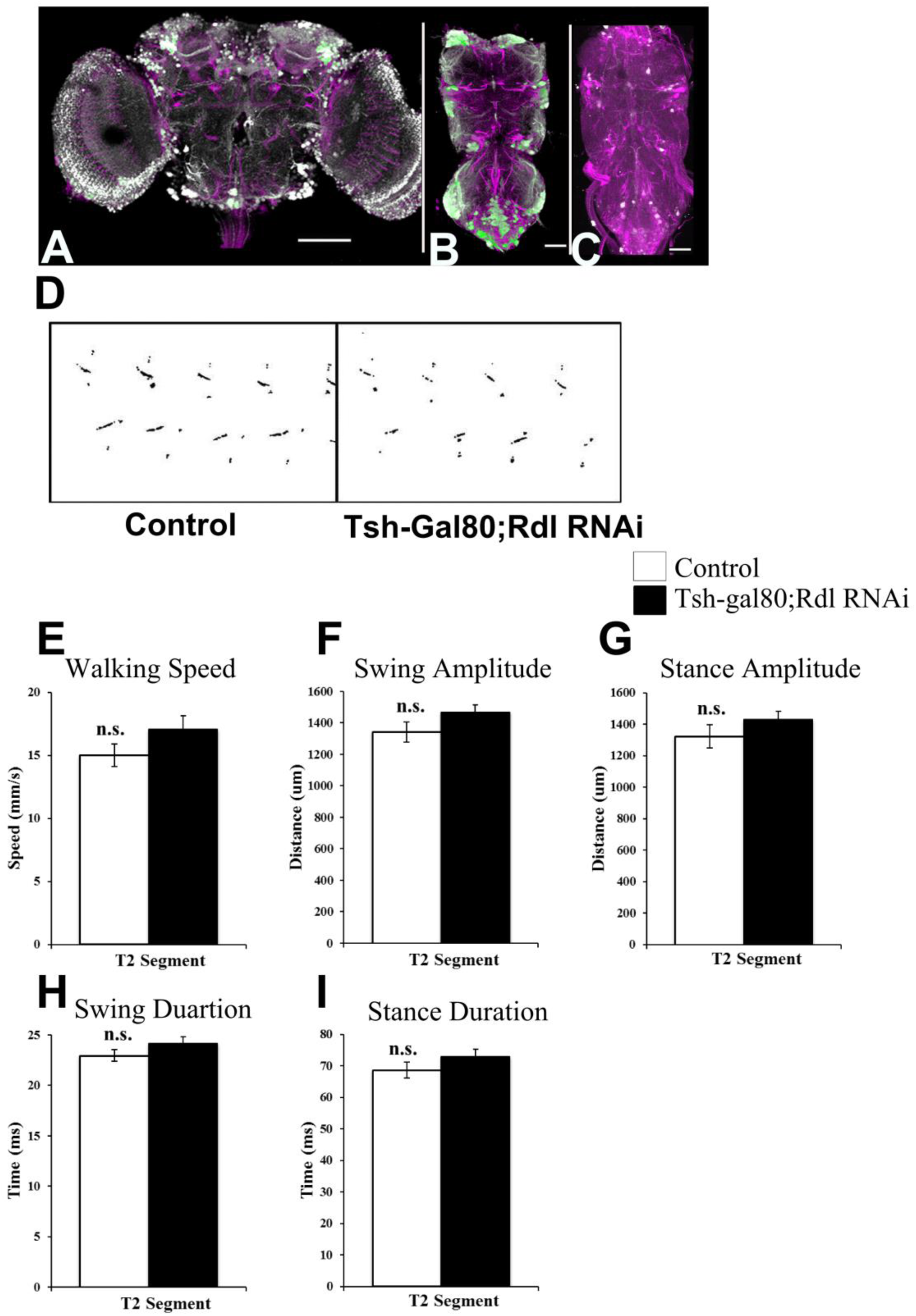
Analysis of Rdl knockdown in Tsh-Gal80 background confirms leg motoneurons as sites of premotor inhibition. (A) OK371 Gal4 driven UAS-eGFP, which is expressed in motoneurons, is also expressed in numerous interneurons of the central brain and optic lobe (Scale bar 100 μm) (B) OK371 Gal4 driven UAS-eGFP expression labels motoneurons in VNC. (Scale bar 50 μm) (C) Tsh-Gal80 inhibits the OK371Gal4 UAS-eGFP expression system in all VNC neurons including motoneurons. N=7. (Scale bar 50 μm) (D) Behavioural analysis of OK371 driven Rdl Knockdown in a Tsh-Gal80 background results in normal walking behaviour as revealed in soot print images. (E) Rdl knockdown on walking behaviour in Tsh-Gal80 background shows no change in walking speed as compared to controls. In this and all subsequent figures left bar indicates control, right bar indicates knockdown. (F-I) Amplitudes and durations of swing and stance are not affected in Rdl receptor knockdown animals using OK371 GAL4 in Tsh-Gal80 as compared to wildtype controls. Controls N=30, Experiment N=30. All the bar represents mean ± SEM (^∗∗∗^: P <= 0.001; ^∗∗^: P <= 0.01; ^∗^: P <= 0.05.). Student’s t-test was performed for all statistical analysis.

To investigate this, we repeated the OK371Gal4 driven Rdl knockdown experiments in a Tsh-Gal80 background. Tsh-Gal80 does not affect the Gal4/UAS system in the central brain (37). However, it inhibits the Gal4/UAS system in all of the neurons of VNC, including motoneurons, as can be seen by using OK371Gal4 to drive a UAS-GFP reporter in a Tsh-Gal80 background (Fig 7B).

Soot print assays reveal that, the walking pattern observed for the OK371 targeted Rdl knockdown animals in a Tsh-Gal80 background corresponds to that of wild type controls (Figure 7D); no walking deficits were apparent in the soot print assay. Quantitative characterization of these walking phenotypes using video analysis also showed that walking speeds (Fig 7E) see **supplementary movie 5**, leg swing and stance phases in these flies were not significantly different as compared to wildtype controls (Fig.7F-I). Thus, no effects on walking were observed when OK371 targeted knockdown is limited to neurons in the central brain. (Data shown here is for T2 leg segment as a representative of other leg segments)

This finding strongly supports the notion that the behavioral phenotypes observed in OK371-Gal4 driven Rdl knockdown experiments is specifically due to the reduction of GABAergic inhibition in leg motoneurons and not in brain interneurons.

## DISCUSSION

In this report, we investigate the role of inhibitory premotor input to leg motoneurons in the walking behavior of freely moving *Drosophila* by suppression of GABAergic inhibitory input to leg motoneurons. Our findings indicate that the reduction of inhibitory GABAergic input to leg motoneurons caused by targeted Rdl down regulation has marked effects on walking behavior. Thus, walking speed was markedly slower as compared to controls and, correlated with this, the amplitudes of the leg protraction (swing) and retraction (stance) phases were significantly smaller and the durations of protraction (swing) and retraction (stance) phases were significantly higher than in controls. Moreover, the concurrency state, in which three legs swing together in a tripod gait, was proportionally reduced as compared to controls. These prominent effects on walking parameters are similar regardless of whether Rdl down regulation occurs throughout development or whether it is restricted to the adult stage. Taken together, these findings reveal a prominent albeit highly specific role of GABAergic premotor inhibitory input to leg motoneurons in the control of normal walking behavior.

The insight into the role of premotor inhibition in walking control reported here is the result of two key experimental methodologies. The first is the highly specific genetic access to identified neuronal populations that can now be attained in the *Drosophila* model system. This is made possible through remarkable targeted expression systems, which together with the availability of libraries of genetically encoded drivers and reporters for molecular manipulation, make it possible to selectively up or down regulate gene expression in highly specific neuronal populations in intact and freely behaving animals (11,13,38). In this study, we have used the Gal4/UAS expression system to achieve targeted genetic access to leg motoneurons and downregulate GABAergic premotor input to these motoneurons by Rdl RNAi expression. In addition we have utilized different forms of the Gal80 repressor to limit Gal4/UAS targeted expression to adult stages or to specific regions of the central nervous system and, hence, refine the spatiotemporal specificity of the resulting genetic access to motoneurons. Given the wealth of Gal4 drivers and UAS-RNAi reporter currently available, it will be possible to use similar transgenic technology to manipulate other inhibitory and excitatory neurotransmitter receptors in future studies of interneuronal components of the walking circuitry.

The second key methods is the development of an advanced automated high-speed video recording and analysis technique that makes it possible to record the protraction (swing) and retraction (stance) phases at high spatiotemporal resolution for each leg in freely walking flies. With this technique, a quantitative assessment of leg motion parameters in freely walking animals can be carried out which can reveal subtle differences in amplitude and phase of movements of individual legs. This quantitative assessment has been critical for uncovering the role of premotor inhibition in walking behavior. Indeed, since the overall leg coordination during walking is unaffected by the reduction of GABAergic input to leg motoneurons, a more conventional qualitative behavioral analysis is unlikely to discern differences in walking between experimental and control animals. Recently, comparable high resolution recording and analysis methods have been used to quantify leg movement parameters in freely walking flies (24,25,39). These studies have provided important information on walking speed, interleg coordination and other locomotor parameters and have also documented a role of sensory proprioceptive input to step precision during walking. The fact that these methods for high resolution recording and analysis are currently available for studying leg movement parameters in freely walking flies should accelerate our understanding of walking behavior and of the neuronal circuitry involved in its control in *Drosophila*.

Given the prominent role of inhibitory premotor input to leg motoneurons reported here, it will now be important to identify and genetically access the premotor interneurons that provide this inhibitory input. While, there is currently little information on the identity of the premotor interneurons that control the activity of leg motoneurons in adult flies, insight into inhibitory premotor interneurons has recently been obtained in larval stages. Thus, in *Drosophila* larva, a set of inhibitory local interneurons termed PMSI neurons have been identified, which control the speed of axial locomotion by limiting the burst duration of motoneurons involved in peristaltic locomotion (32). Moreover, a second set of inhibitory premotor interneurons called GVLI neurons have been reported which may be part of a feedback inhibition system involved in terminating each of the waves of motor activity that underlie larval peristalsis (33). Finally, a pair of segmentally repeated GABAergic interneurons termed GDL neurons have been identified which are necessary for the coordinated propagation of peristaltic motor waves during both forward and backward crawling movements of larvae (34). Whether or not these inhibitory premotor interneurons persist into the adult stage and act in the control of walking behavior is not known.

In general terms there are fundamental similarities in the principle mechanisms of locomotion in insects and vertebrates (e.g. 3). These mechanistic similarities might also reflect similar motor circuit properties. For example, much like the PMSI neurons in *Drosophila*, which control the speed of locomotion by limiting motoneuron burst duration, the premotor V1 spinal interneurons in mammals are involved in the regulation of leg motoneuron burst and step cycle duration and thus is also likely control the speed of walking movements (40). Hence, a characterization of the behavioral effects of inhibitory input to leg motoneurons in *Drosophila*, notably in freely walking flies, is likely to provide useful comparative information for understanding the functional role, and possibly the evolutionary origin, of premotor inhibition in vertebrate locomotory circuitry.

## MATERIAL AND METHODS

### Fly strains used

OK371-Gal4, UAS-dicer; OK371-Gal4, TubGal80ts, UAS-dicer; OK371 Gal4, UAS-mCD8GFP; VGN 1-Intron Reg-3 Gal4. UAS-Rdl i8-10G RNAi from Ron Davis (41); UAS-Trip RNAi – 31622 (Bloomington). Tsh-Gal80,UAS-eGFp(37); Tsh-Gal80, UAS-Rdl RNAi.

### Generation of VGN 1-Intron Reg-3 Gal4 transgenic flies

DVGlut regulatory region of a 640 bp, corresponding to genome coordinates 2L:2403206-2403845 (release 6.16) was PCR amplified from wild type *Drosophila* melanogaster canton-s and cloned into restriction endonuclease enzyme sites, EcoR1 and BamH1 of pPTGal-attB vector (42) to generate VGN-Intron region3 Gal4.

### Behavioural analysis procedure

#### Soot assay procedure

Soot assay analysis was done as previously described (17)

### Video recording procedure

Recording walking behavior of freely moving flies: In order to record leg movements of freely walking flies at high spatio-temporal resolution, flies were placed in a transparent plexiglass arena and their locomotor behavior was recorded by an overhead video camera at 200 frames/s (Fig. 1A).The arena had a square-shaped flat central bottom surrounded by two opposing inclined surfaces and had Teflon coated walls to discourage climbing. Single fly with wings clipped were introduced into the arena and preferentially walked across the flat part of its bottom. The recording video camera was fitted with a lens of 10x magnification. Illumination was from below resulting in high contrast images (Fig 1B). An automated recognition procedure was employed to extract the leg movements of the freely walking fly from the recorded video images; only linear walking sequences were selected. For this, thresholding based on light intensity differences was used to demarcate the fly torso, watershed filtering was employed to delimit head, thorax and abdomen and computation of a body axis orientation vector based on these three body parts was carried out (Fig. 1C). Subsequently, based on the dynamic changes of this vector, the fly’s motion was transformed from the camera’s frame of reference to an inertial reference frame (fly’s frame of reference) (Fig. 1D). This provided a stable image of the fly suitable for auto-detection and analysis of leg movements. 3-4 day old flies starved for 2-3 hours were used for recording.

### Immunohistochemistry and Confocal Microscopy

Adult brain and thoracic ganglia were dissected in phosphate buffer saline (pH 7.8) (PBS) and fixed in 4% buffered formaldehyde for 45 min at 4°C. Staining procedures was performed as previously described (17). Imaging of adult brain and thoracic ganglia were performed using Olympus FV 1000 confocal microscope at 1 micron intervals. The imported Z stack images were processed using FIJI and Adobe photoshop for further adjustments in brightness and contrast.

### Antibodies used

#### Primary antibodies

Chicken pAb a-GFP (1:1000, Abcam), Rabbit (Rb) a-GFP (1:4000, Abcam), mouse (ms) a-Neuroglian (BP104, 1:40;DSHB).

#### Secondary antibodies

Alexa fluor-488 and 647 (1:400) from Invitrogen were used in all staining procedures.

## ACKNOWLEDGEMENTS

We are thankful to Ron-Davis, the VDRC, the Bloomington Stock Centre and the NCBS fly facility for generously providing fly stocks and other reagents. We acknowledge Avinash, Akila Sridar and Swati Krishnan for their contribution in preliminary soot assay experiments. We are greatly thankful to Sanjay Sane and Vatsala Thirumalai for their valuable suggestions and comments. We thank Central imaging and flow facilities for the Olympus FV1000 microscopes at NCBS. This work was supported by a J.C.Bose Fellowship of K.VijayRagahavan.

